# Inheritance of Repressed Chromatin Domains during S-phase Requires the Histone Chaperone NPM1

**DOI:** 10.1101/2021.08.31.458436

**Authors:** Thelma M. Escobar, Jia-Ray Yu, Sanxiong Liu, Kimberly Lucero, Nikita Vasilyev, Evgeny Nudler, Danny Reinberg

## Abstract

The epigenetic process safeguards cell identity during cell division through the inheritance of appropriate gene expression profiles. We demonstrated previously that parental nucleosomes are inherited by the same chromatin domains during DNA replication only in the case of repressed chromatin. We now show that this specificity is conveyed by NPM1, a histone H3/H4 chaperone. Proteomic analyses of late S-phase chromatin revealed NPM1 in association with both H3K27me3, an integral component of facultative heterochromatin and MCM2, an integral component of the DNA replication machinery; moreover NPM1 interacts directly with PRC2 and with MCM2. Given that NPM1 is essential, the inheritance of repressed chromatin domains was examined anew using mESCs expressing an auxin-degradable version of endogenous NPM1. Upon NPM1 degradation, cells accumulated in S-phase of the cell-cycle and parental nucleosome inheritance from repressed chromatin domains was markedly compromised. Appropriate inheritance required the NPM1 acidic patches that function in chaperone activity, pointing to NPM1 being integral to the epigenetic process.

**One-Sentence Summary:** The histone H3/H4 chaperone, NPM1, fosters epigenetic inheritance from parental repressed chromatin during DNA replication.

## Main Text

A hallmark of the epigenetic process entails the regulated inheritance of a sufficient platform of gene expression patterns from parental cells such that the original gene expression profile is fully recapitulated in progeny cells; bypassing the re-establishment *de novo* of a particular cell identity. The structure of chromatin domains can provide such a platform, exhibiting features that either promote chromatin accessibility to the transcription machinery or foster its compaction. Whether or not chromatin domains are heritable was recently resolved (*1–3*). A novel CRISPR-Cas9 Biotinylation System revealed the profile of parental nucleosome segregation during DNA replication with single locus specificity: parental nucleosomes are indeed inherited to the same chromatin domains during S-phase, but only in the case of repressed and not active chromatin domains (*1*). Importantly, transcription activation of a previously repressed locus leads to the dispersal of parental nucleosomes, thereby thwarting inheritance (*1*). This finding points to features inherent to the repressed chromatin state being epigenetic.

Facultative heterochromatin comprises di- and tri-methylated lysine 27 of histone H3 (H3K27me2/me3). The multi-subunit Polycomb Repressive Complex 2 (PRC2) contains a subset of the Polycomb group of proteins and is the sole enzyme responsible for all states of H3K27 methylation (*4–8*). Notably, the EED subunit of PRC2 recognizes the product of PRC2 catalysis, H3K27me3, resulting in an allosteric activation of PRC2 and stimulation of its histone methyltransferase activity conveyed by its EZH2 subunit (*9*). This feed-forward mechanism accounts for the formation of extensive, repressive facultative heterochromatin domains (*4, 10*). As well, given that only repressed chromatin domains are inherited, we postulate that this feed-forward mechanism can account for the full restoration of repressive chromatin domains upon DNA replication: H3K27me3 within locally segregated parental nucleosomes provides the allosteric activator that stimulates PRC2 catalysis of the tri-methyl modification on newly incorporated naïve nucleosomes (Fig. 1A). Yet, these findings and the ensuing reasoning beg the question as to what process provides specificity such that parental nucleosome inheritance is limited to repressed chromatin domains. To address this aspect of the epigenetic process, we first investigated proteins associated with late S-phase replicating, facultative heterochromatin.

**Figure 1.**
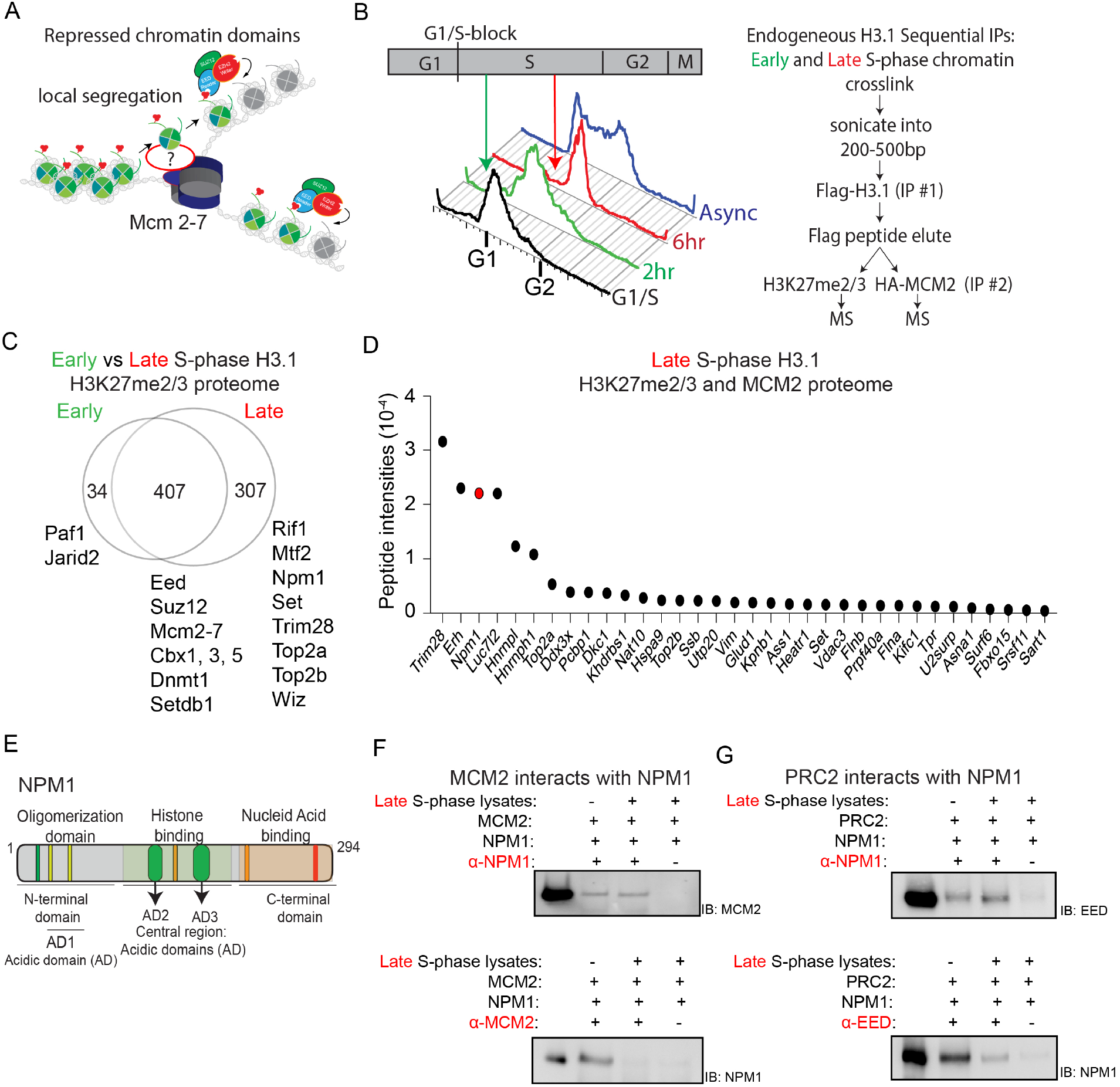
H3.1-proteome analysis identifies NPM1 in late H3K27me2/3-replicating chromatin domains. **(A)** Schematic illustration of the restoration of fully repressed chromatin domains across the DNA replication fork (MCM2-7). Parental nucleosomes (green) are shown segregating randomly to the leading and lagging DNA strands within the same repressive chromatin domains. PRC2 recognizes H3K27me3 (red triplet) within parental nucleosomes through its EED subunit resulting in its allosteric activation and propagation of H3K27me3 to naïve nucleosomes (grey). **(B)** Experimental set-up for chromatin immunoprecipitations from mESCs followed by mass spectrometry (ChIP-MS) to identify H3.1-associated proteins in late S-phase. mESCs synchronized at G1/S by a double thymidine block were released for either 2 hr (Early S-phase) or 6 hr (Late S-phase) and chromatin isolated. Sequential ChIPs were performed to isolate proteins associated with H3.1 followed by those associated additionally with H3K27me2/me3 or HA-MCM2. **(C)** Unique polypeptides identified from H3.1 and H3K27me2/3 proteome analysis within early and late S-phase reproduced from four independent experiments. **(D)** The 307 polypeptides identified in “Late” from **(C)** were crossed-referenced with the Late MCM2-replicative proteome (two independent experiments) to identify the H3.1-proteome present in IPs from both H3K27me2/me3 and HA-MCM2. Protein intensities were calculated by the sum of peptide intensities extracted from MS1 spectra minus the polypeptides obtained from IPs of untagged-H3.1 control. **(E)** Schematic showing NPM1 protein domains with its N-terminus comprising an oligomerization domain and an acidic domain (AD1) and its central region containing two acidic domains (AD2 and AD3) that exhibit histone binding and chaperone activities, and its C-terminal domain comprising nucleic acid binding activity and a nucleolar localization sequence. **(F, G)** Reciprocal immunoprecipitations of recombinant NPM1 and either recombinant MCM2 **(F)** or core PRC2 **(G)** with and without incubation with late S-phase lysates, as indicated. NPM1 interacts directly with MCM2 **(F)** and with core PRC2 **(G)**.

As euchromatic regions are replicated at early times of S-phase and heterochromatin is replicated later (*11*), we compared a proteomic analysis of chromatin containing H3K27me2/me3 at early and late times of S-phase. Mouse embryonic stem cells (mESCs) expressing endogenous, FLAG-BAP-tagged versions of four H3.1 alleles (Figs. S1A and B, (*3*)) were infected with lentivirus expressing an HA-tagged version of MCM2 (Fig. S1C), an integral component of the DNA replication machinery that makes the first contact with parental nucleosomes (*12, 13*). After synchronization at the G1/S boundary using a double thymidine block, the cells were released into S-phase for 2 or 6 hr (Early or Late S-phase, respectively). Chromatin specific to Early or Late S-phase was isolated, cross-linked, and 200-500 bp fragments were subjected first to FLAG-H3.1 IP with the resulting eluate being halved and subjected to IP against either H3K27me2/me3 or HA-MCM2 (Fig. 1B). Mass spectrometric analyses identified specific proteins associated with both H3.1 and H3K27me2/me3 in Early versus Late S-phase cells (Fig. 1C). PRC2 association was detected in both cases as evidenced by the presence of its EED and SUZ12 subunits, as were components of the DNA replication machinery, MCM2-7. Of note, the H3/H4 histone chaperone, NPM1, was specific to the Late S-phase H3.1/H3K27me2/me3 proteome (Fig. 1D). Interestingly, our previous proteomic analysis involving PRC2 revealed its association with NPM1 (*10*). Importantly, amongst the proteins identified as being common to both H3.1/H3K37me2/me3 and H3.1/MCM2 proteomes from Late S-phase, NPM1 was one of the prominent candidates (Fig. 1D).

We next examined whether NPM1 interacts with PRC2 and/or MCM2 using recombinant versions of NPM1 and MCM2 (Fig. S1D), and the purified core PRC2. Reciprocal IPs demonstrated that NPM1 interacted directly with MCM2 and that this interaction did not require other proteins present in Late S-phase extract, the addition of which appeared instead to be interfering (Fig. 1F). Similarly, reciprocal IPs demonstrated a direct interaction between NPM1 and PRC2, and the addition of Late S-phase extract in this case was ineffectual (Fig. 1G). Thus, NPM1 interacted directly with major activities involved in either DNA replication (MCM2) or facultative heterochromatin formation (PRC2).

Amongst its many reported biological functions (*14*), NPM1 interaction with histones has implicated this protein in several chromatin-based processes, including DNA replication and repair, transcription, and chromatin remodeling (*12, 15*). The distinct NPM1 protein domains include a central region with acidic patches (Fig. 1E), which exhibits histone binding activity with a strong preference for histone H3/H4 tetramers, relative to histone H2A/H2B dimers (*16*). These acidic regions are critical for the histone H3/H4 chaperone activity of NPM1 *in vitro* (*16*) and *in vivo* (see below, Fig. S2). Given its reported H3/H4 chaperone activity *in vitro* (*16, 17*), and our findings here that NPM1 associates with Late S-phase chromatin and directly interacts with both MCM2 and PRC2, we investigated the possibility that NPM1 facilitates repressed chromatin domain inheritance.

As NPM1 is required for cell viability (*18*), we employed the auxin-inducible-degradation (AID) system (*19, 20*) to deplete cellular NPM1 upon auxin addition. The endogenous NPM1 gene in mESCs having four FLAG-BAP tagged versions of histone H3.1 used above (Fig. S1A) was engineered to express an AID-tag at its N-terminus using CRISPR-Cas9 technology (Fig. 2A, left and see below). Indeed, AID-NPM1 was undetectable by western blot (WB) at 6 hr of auxin treatment (Fig. 2A, right). To ascertain a time period during which auxin addition would be sufficient to deplete AID-NPM1 without affecting cell survival, but also be tenable for analyzing parental nucleosome inheritance as preformed previously (*1*), we first analyzed cells treated for 24 and 48 hr with auxin. Protracted auxin treatment gave rise to severe defects in cell proliferation evident after 24 hr (Fig. 2B). At 24 hr, the AID-NPM1 cells already exhibited an abnormal blockage at G1, relative to untreated cells (Fig. 2C). The salient features associated with facultative heterochromatin were gauged as a function of time after auxin addition by WB. While AID-NPM1 was undetectable by 6 hr of auxin treatment, auxin treatment for up to 24 hr was ineffectual with respect to levels of the core PRC2 subunits: SUZ12, EZH2, and EED (data not shown).

**Figure 2.**
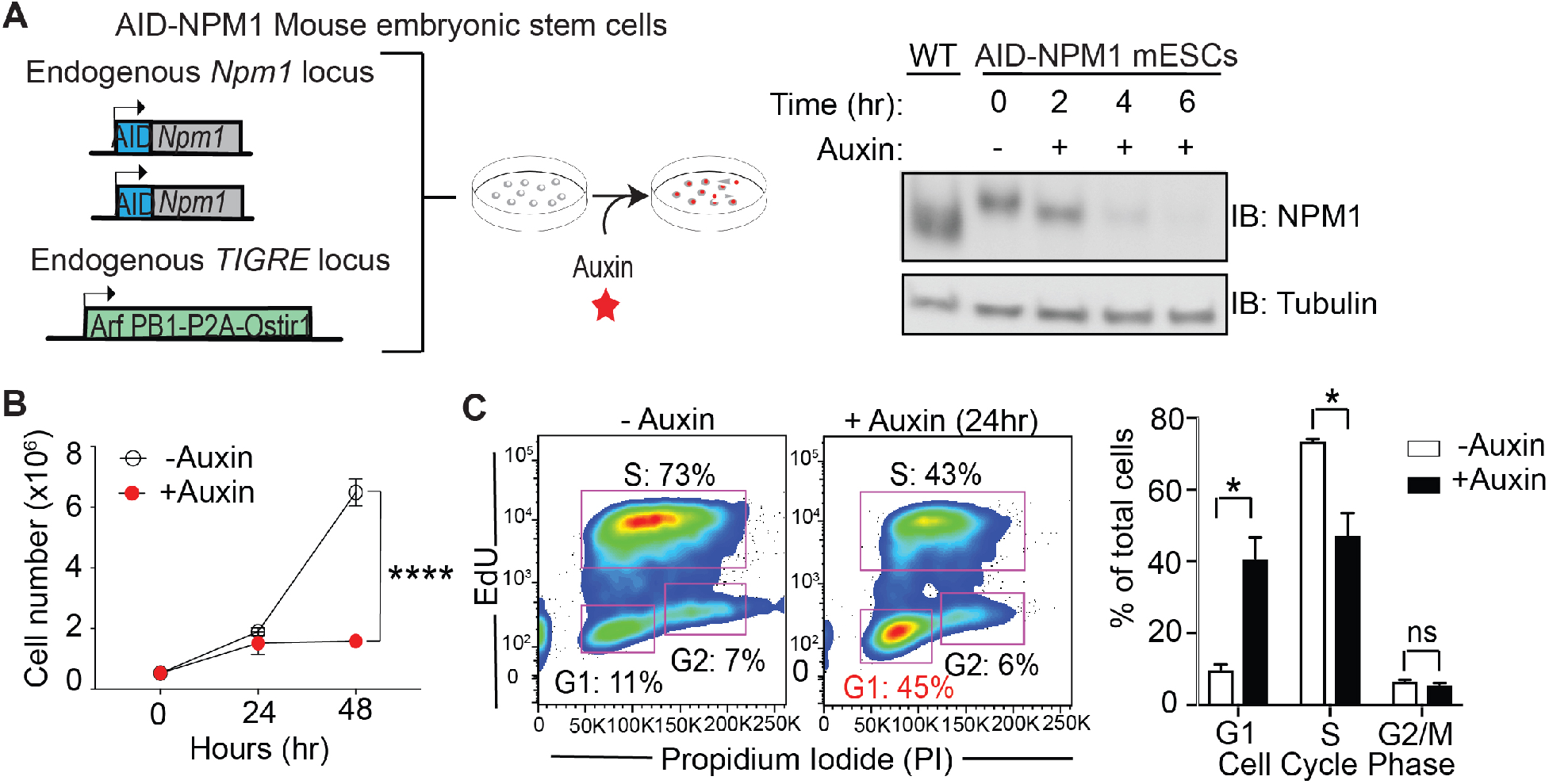
Cell-cycle effects accompanying NPM1 depletion. (**A**) AID-NPM1 mESCs were generated using CRISPR knock-in such that an AID tag was appended to the 5′-end of the *Npm1* gene and the auxin-binding receptor, Oryza sativa TIR (Tir1), to the TIGRE locus (left). Western blot analysis of whole cell extracts following the addition of auxin to AID-NPM1 mESCs in a time-dependent manner (0−, 2−, 4−, and 6-hr time points) showing that NPM1 is depleted within 6 hr (right). **(B)** Cell proliferation analysis of AID-NPM1 mESCs in the presence and absence of auxin over a 48-hr time period. Statistical significance was determined by 2-way ANOVA (****p<0.0001). **(C)** Cell cycle analysis by flow cytometry of AID-NPM1 mESCs in G1−, S-, and G2/M-phase based on the analysis of Propidium Iodide (DNA content) and EdU incorporation, in the presence and absence of auxin. The percentage of cells in each phase is shown on the right with significance determined by Student t-test (*p<0.05).

These findings suggested that a 12 hr auxin treatment would not only be sufficient for AID-NPM1 depletion, but also satisfy the 12 hr time frame previously established for examining parental nucleosome inheritance after release into S-phase ((*1*) and see below). Thus, we next examined the phenotype of these AID-NPM1 cells that were blocked at the G1/S boundary and then released into S-phase for 12 hr, with and without auxin treatment (Fig. 3A, top). As expected, NPM1 was depleted within 6 hr (Fig. 3A, bottom). The cell cycle profile after release into S-phase for 6 hr was similar, without and with auxin treatment (Fig. 3B, top middle and right panels, respectively), while cells released into S-phase for 12 hr showed evidence of a G1 blockage in the subsequent cell cycle in the case of auxin-treated, relative to untreated cells (Figs. 3B, bottom right and middle panels, respectively, and 3C). This perturbation of the cell cycle profile was accompanied by the occurrence of differentially expressed genes (DEG), reflective of aberrant up- and down-regulated gene expression (Fig. 3D). Notably, PRC2-target genes were more enriched in the up-regulated dataset (Fig. 3E), suggesting that NPM1 depletion alters gene repression. A GO analysis indicated that the top 5 de-repressed genes were related to development (Fig. 3F). Of note, a naturally occurring, heterozygous mutated *NPM1* allele (NPM1c+) exhibits abnormal cytoplasmic retention and is associated with ~35% of Acute Myelogenous Leukemia (AML) with normal karyotype (*21*). Previous studies demonstrated that *NPM1c+* AML exhibit de-repression of specific *HOXA and HOXB* loci (*22*). In accordance, the AID-NPM1 cells exhibited increased transcription at the *HoxA9*, *HoxB4* and also at the *Gata2* loci as a function of auxin treatment for 12 hr after S-phase release, as evidenced by RNA-seq (Fig. 3G).

**Figure 3.**
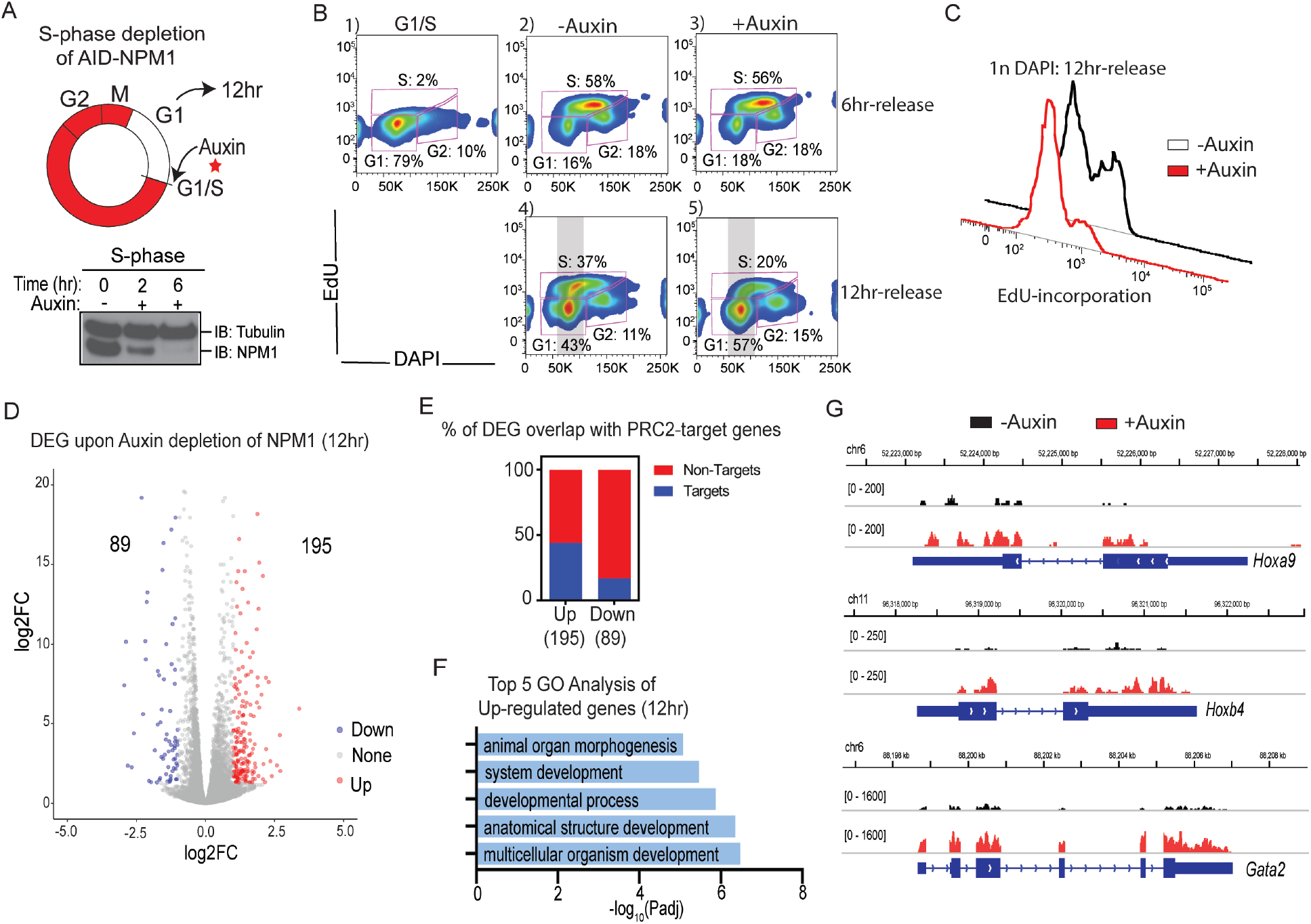
Altered gene expression upon depletion of NPM1. **(A)** Schematic showing the protocol for S-phase specific addition of auxin to AID-NPM1 cells (top). Cells were synchronized at G1/S with a double thymidine block and auxin was added upon washing and releasing cells into S-phase. WB analysis shows depletion of NPM1 within a 6-hr release into S-phase (bottom). For further analysis as in **(B)**, cells were also harvested within the next G1-phase (12-hr release). **(B)** Cell cycle analysis by flow cytometry of AID-NPM1 cells in the presence and absence of auxin during a 6-hr and 12-hr release from the G1/S-block. G1−, S-, and G2/M-phase were designated based on DAP1 analysis and EdU incorporation. **(C)** Analysis of EdU incorporation by AID-NPM1 cells upon a 12-hr release in the presence and absence of auxin, taken from the DAPI stain within the gray area indicated in **(B)**. **(D)** Differentially expressed genes (DEG) identified by RNA-seq analysis of AID-NPM1 cells upon a 12-hr release in the presence and absence of auxin (two replicates each). **(E)** Percentage of DEGs from **(D)** that overlaps with PRC2/H3K27me3 ChIP-seq targets in mESC. **(F)** The Gene Ontology (GO) term enrichment of those genes that were de-repressed in the presence of auxin as identified in **(D)**. **(G)** Genome-browser screenshots of RNA-seq at *Hoxa9*, *Hoxb4*, and *Gata2* loci depicting results as normalized coverage tracks upon a 12-hr treatment with (red) and without (black) auxin in AID-NPM1 mESCs. Data represents one replicate of two.

To examine if NPM1 fosters epigenetic inheritance and in particular, through H3K27me3-modified nucleosomes, we re-visited the CRISPR-Cas9 Biotinylation system that captured the inheritance of biotinylated parental nucleosomes specifically from repressed loci during DNA replication at single locus resolution (*1*), but now in the context of NPM1 depletion. We examined anew two such loci, *Gata2* and *Gata6*, whose expression is inducible and which lost parental nucleosome inheritance upon activation with retinoic acid (*1*). We re-built the CRISPR-Cas9 Biotinylation System (Fig. 4A) in the AID-NPM1 cell line that expresses four FLAG-BAP (biotin acceptor protein)-tagged H3.1 alleles. We stably expressed a doxycycline-inducible version of dCas9 that is fused to BirA that biotinylates BAP (dCas9-BirA), as used previously. In lieu of the FKBP degron used previously, dCas9-BirA was now fused to an AID tag (dCas9-BirA-AID) that similarly ensures its presence is limited to the G1-phase and its absence in S-phase. Using this system, we gauged parental nucleosome inheritance as a function of the presence of NPM1 using stably expressed sgRNAs that specifically target dCas9-BirA-AID to the *Gata2* or *Gata6*.

**Figure 4.**
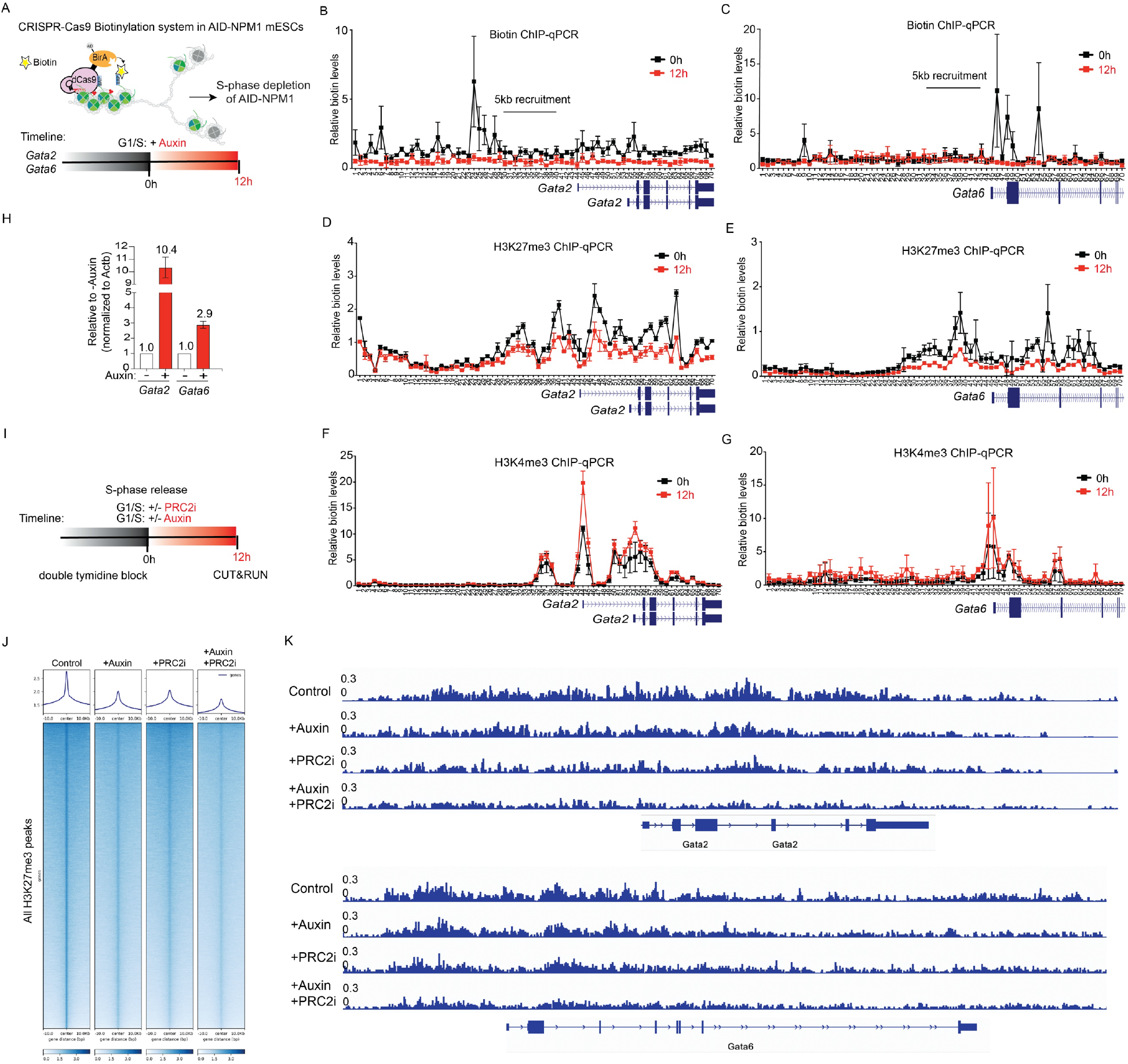
Depletion of NPM1 thwarts inheritance of repressed chromatin domains. **(A)** Schematic of the previously reported CRISPR-Cas9 Biotinylation System that captured the inheritance of repressed, but not active, chromatin domains during S-phase, in this case using AID-NPM1 cells treated or untreated with auxin. As previously described (Escobar et al. 2019), G1/S blocked AID-NPM1 cells received a 6-hr pulse of Dox to induce expression of an engineered dCas9-BirA-AID fusion protein, exogenous biotin to mark nucleosomes, and sgRNAs for targeting BirA to the *Gata2* or *Gata6* loci. Cells were then released into S phase for 12 hr as previously described, but now in the presence or absence of auxin. Cells were then processed for **(B, C**) native mononucleosomal biotin ChIP-qPCR for *Gata2* **(B)** or *Gata6* **(C)** loci, **(D, E)** H3K27me3 ChIP-qPCR for *Gata2* **(D)** or *Gata6* **(E)** loci, and **(F, G)** H3K4me3 ChIP-qPCR for *Gata2* **(F)** or *Gata6* **(G)** loci. **(H)** Relative mRNA expression of *Gata2* and *Gata6* genes in AID-NPM1 cells in the absence and presence of auxin treatment for the 12-hr period. Data was normalized to Actin expression and (−) auxin control. **(I)** A depiction of the experimental flow for single or double treatment with auxin or/and PRC2 inhibitor (GSK126), followed by CUT&RUN. **(J)** Heatmaps of H3K27me3 CUT&RUN using AID-NPM1 cells treated under the conditions described in **(I)**. Top, line plots of relative quantifications based on averaged intensities of all H3K27me3 peaks detected. Bottom, all H3K27me3 peaks centered by max peak intensity within a +/−10 kb window. **(K)** Representative tracks of the *Gata2* and *Gata6* loci from the H3K27me3 CUT&RUN experiments described in **(I)** and **(J)**.

G1/S arrested AID-NPM1 cells were treated with limited doxycycline to induce dCas9-BirA-AID expression and biotin to label the appropriate FLAG-BAP-H3.1 for 6 hr, as performed previously (*1*). The cells were then released into S-phase for 12 hr as performed previously, but this time as a function of the presence of auxin that targets both AID-NPM1 and dCas9-BirA-AID. ChIP-qPCR for biotin demonstrated that the parental nucleosome inheritance evident at both the *Gata2* and *Gata6* loci in the case of untreated cells was lost in the case of auxin-treated cells that were depleted of NPM1, (Fig. 4B and C, respectively), strongly suggesting that NPM1 is required for parental nucleosome inheritance from repressed loci. Yet, while the biotin-tagged nucleosomes dispersed from these repressed loci upon DNA replication in the absence of NPM1, nucleosomes comprising the repressive histone modification, H3K27me3, were clearly evident, albeit at a lower level (Fig. 4D and E). Notably, these developmentally regulated genes are inherently bivalent, comprising both H3K27me3 and H3K4me3 (*23–26*). Indeed, an increased presence of H3K4me3 was evident at both loci, under these conditions (Fig. 4F and G). Given that PRC2 associates with replicating DNA (*27–29*), we reasoned that the levels of H3K27me3 detectable in the absence of NPM1 might arise *de novo* from PRC2*-*mediated catalysis of H3K27me3 on randomly deposited naïve nucleosomes.

To examine the contribution of *de novo* H3K27me3 catalysis by PRC2 upon release from S-phase, we treated the synchronized AID-NPM1 cells with either auxin, or the PRC2 inhibitor (PRC2i), GSK126 that targets EZH2, or both (Fig. 4I). We used a high dose of GSK126 (5 μM) in order to achieve an efficient and acute inhibition of PRC2. As expected, treatment with auxin or PRC2i resulted in a moderate, ~50% reduction of global H3K27me3 compared to control cells (shown by line plots and heatmaps from H3K27me3 CUT&RUN experiments, Fig. 4J). However, cells co-treated with auxin and PRC2i exhibited a considerably greater reduction in H3K27me3 deposition after exiting S-phase (12 hr). This effect was not only apparent at the *Gata2* and *Gata6* loci (Fig 4K), but also genome-wide (Fig. 4J), consistent with NPM1 being key to histone re-deposition at repressive loci.

NPM1 is a demonstrated chaperone in nucleosome assembly, with its central region containing two acidic patches/domains (AD2 and AD3) that bind to histones and are required for its chaperone activity (Figure 1E, (*16, 30*)). Thus, we probed AD2, AD3, and AD2+3 compound mutants to ascertain whether the chaperone activity of NPM1 is involved in the inheritance of repressed chromatin domains. We performed rescue experiments by ectopically expressing HA-tagged versions of NPM1, either wild type or mutant in the indicated acidic patch(es), in auxin-treated AID-NPM1 cells (Fig. S2A). Biotin-labeling through dCas9-BirA-AID was targeted to the representative *Gata2* locus. Although the polyclonal NPM1 antibody does not detect the NPM1 mutants efficiently, the expression of ectopically-expressed NPM1 candidates was comparable in all cases as confirmed by HA antibody (Fig. S2A). Biotin-ChIP-qPCR indicated that while auxin-resistant, wild type NPM1 rescued the deficiency of histone re-deposition, each of the NPM1 chaperone mutants failed to rescue this inheritance (Fig. S2B), underscoring that NPM1 chaperone activity is required for this feature of the epigenetic process.

## Acknowledgments

The authors are grateful to Dr. Lynne Vales for active discussions and help in writing the paper, to Dr. Alejandra Loyola for active discussions, and to Deborah Hernandez for her impeccable technical assistance. TE was supported by an 3R01CA199652-14S1 grant, J-R Y was supported by the American Cancer Society (PF-17-035-01), SL was supported by the Howard Hughes Medical Institute. All studies were supported by a grant from the NCI (9R-1CA199652-13A1) as well as by HHMI.

## Authors contributions

TE directed some of the experiments, performed most of the experiments, and wrote most of the Figure Legends and Materials and Methods. J-R Y performed and analyzed experiments described in Figure 4, J & K and Supplemental Figure 2 and wrote accompanying legends. SL performed bioinformatic analysis and helped with the interpretation of sequencing results. NV performed LC-MS and normalization of the raw data. DR directed the experiments and oversaw the manuscript.

## Data and materials availability

The accession numbers for the raw data FASTQ files and processed files for all sequencing data will be available on GEO upon publication. All plasmids are available upon request and modified KH2 mES cell lines are available through a materials transfer agreement.

## Supplementary Materials

### Materials and Methods

#### Cell line generation

KH2 mouse ESCs used in this study were grown in DMEM supplemented with 15% FBS L-glutamine, penicillin/streptomycin, non-essential amino acids, 0.1 mM β mercaptoethanol, LIF, and 2i inhibitors, which include 1 μM MEK1/2 inhibitor (PD0325901) and 3 μM GSK3 inhibitor (CHIR99021) on 0.1% gelatin coated plates. 293T cells were used for lentiviral production and grown in DMEM supplemented with 10% FBS, L-glutamine and penicillin/streptomycin.

Targeting of mESCs to endogenously express H3.1 genes comprising FLAG-Biotin Acceptor Peptide (BAP) at the N-terminus was described previously ((*1*); cell line 30-55-15), with the genotype of the cells used in this study presented in Fig. S1A. Briefly, gene editing of the *Hist1h3* locus to incorporate FLAG-BAP tags was done by using ~750 bp gBlock for each gene, ordered from IDT or Genscript. These gBlocks included: 1) a homology arm corresponding to ~350-450 bp of the H3 promoter area, 2) the FLAG-BAP sequence placed after the start codon (FLAG-BAP sequence: GACTACAAAGACGATGACGACAAGGGCCTGACAAGAATCCTGGAAGCTCAGAAGATCGTGAGAGGAGGCCTCGAG), and 3) a homology arm corresponding to ~200-300 bp of the H3.1 coding sequence. The PAM sequences of the gBlocks were mutated for correct Cas9 digestion of genomic DNA within cells. ESCs were then transfected with 0.5 μg of Cas9-gRNA-BFP plasmid targeting the promoter of H3.1 genes and 0.5 μg of amplified gBlock in Lipofectamine 2000 (Invitrogen) containing 2i media. Transfected cells were FACS sorted and seeded at 20,000 cells per 15 cm plate and 7-10 days later, single ESC clones were selected and plated onto individual wells of a 96-well plate for genotyping. Genomic DNA was harvested via QuickExtract (Epicentre) DNA extraction, and genotyping PCRs were performed using primers surrounding the target site. The PCR products of the FLAG-BAP positive clones were purified and sequenced to verify the presence and correct sequence of a FLAG-BAP-H3 insertion. Primers for gRNA, gBlocks, and genotyping are shown in Table S1.

For targeting of *NPM1*, a gBlock of approximately of 1897 bp was ordered from Genscript that included: 1) a homology arm corresponding to ~850 bp of the *NPM1* promoter, 2) the *IAA17* 71-113 mRNA sequence with a 3x Glycine linker placed after the start codon (AID-3xGly sequence: CCTAAAGATCCAGCCAAACCTCCGGCCAAGGCACAAGTTGTGGGATGGCCACCGGT GAGATCATACCGGAAGAACGTGATGGTTTCCTGCCAAAAATCAAGCGGTGGCCCGGAGGCGGCGGCGTTCGTGGGTGGAGGT), and 3) a homology arm corresponding to ~900 bp of the *NPM1exon1* and *NPM1intron1*. The PAM sequence of the gBlock was mutated to correct Cas9 digestion of the genomic DNA within cells and cells were transfected with 0.5 μg of the Cas9-*NPM1*gRNA-GFP plasmid, FACS sorted, and genotyped as above (Table S1 for gRNA, gBlocks, and genotyping primer sequence). To check for correct in-frame insertion of *IAA17* to the *NPM1* transcript, RNA was purified from homozygous *IAA17-NPM1* genotype mESCs followed by cDNA synthesis. The cDNA was then used to amplify the *IAA17-NPM1* full-length transcript with a Q5/Taq Polymerase mix and subsequently cloned into the TOPO-TA vector. The *IAA17-NPM1* mRNA transcript in the TOPO vector was sequenced using the M13 primers. The mESC clones that showed the correct sequence were selected and whole cell extract taken for western blots to observe the kDA shift increase in NPM1 protein (AID-NPM1). Finally, to obtain an optimized auxin-induced degron system (*20*), these AID-NPM1 mESCs were targeted at the TIGRE locus to contain ubiquitous expression of the auxin-binding receptor, *Oryza sativa TIR* (OsTir1 or TIR) (*20*). TIGRE gRNAs and a GFP-ARF16-PB1-OsTIR gBlock were transfected into AID-NPM1 mESCs and screened for GFP integration and auxin inducibility of AID-NPM1 degradation in GFP+ mESCs. For auxin-induced degradation of AID-NPM1, 2.5 mM auxin was added to the cultures for the time frames indicated in the Figures.

To generate dCas9-BirA-AID expressing stable cell lines used for biotinylation experiments, 2 μg of pINTA-dCas9-BirA-AID plasmid was transfected with Lipofectamine 2000 into FLAG-BAP-H3.1; AID-NPM1; TIR KH2 ESCs. After 2 weeks of 200 μg/ml of Zeocin selection, single colonies were selected and dCas9-BirA-AID inducibility tested with doxycycline. Cells were then transduced with gRNAs tiling the *Gata2* and *Gata6* locus, as described previously (*1*) for downstream biotinylation experiments. For proteomics studies, which required expression of HA-MCM2 in mESCs, HA-MCM2 was inserted into pLVX-EF1a-IRES-ZsGreen1 (see cloning below), lentiviruses made and transduced into FLAG-BAP-H3.1 KH2 mESCs. Lentivirus production in 293T cells and transduction of KH2 mESCs was previously described (*1*).

#### Cell cycle synchronization and analysis

For G1 synchronization experiments, mESCs were plated at 50-60% confluency and pulsed for 18 hr with 8 μM thymidine, followed by a PBS wash and a 7-8 hr release into 2i media. mESCs were then given a second thymidine treatment at a 5-6 μM concentration for 12-13 hr. The G1-block was confirmed by staining DNA with propidium iodide. For cell cycle analysis using EdU labeling, Invitrogen’s Click-iT EdU Cell Proliferation Kit for Imaging 488/564 dye was used. The biotinylation experiments were performed as described previously (*1*), however, for auxin depletion of NPM1 in S-phase, cells were given 2.5 mM auxin immediately after releasing cells from a G1/S-block after the second thymidine treatment.

#### Cloning

To generate pLVX-EF1a-IRES-ZsGreen1-HA-MCM2, mouse MCM2 cDNA (Horizon discovery-Dharmacon) was amplified with Q5 polymerase, using primers containing an HA tag followed by a 5xGly linker synthesized by IDT (see Table S2). PCR product and pLVX-EF1a-IRES-ZsGreen1 vector were digested with Xba1 and BamH1, ligated, and transformed into XL10 Gold competent cells. Colonies were picked and correct cloning confirmed by sequencing. To generate constructs pFastbac1-6xHis-FLAG-TEV-NPM1 and pFastbac1-6xHis-FLAG-TEV-MCM2 and to purify NPM1 and MCM2 protein from SF9 cells, primers were synthesized from IDT containing 6xHis- and FLAG-tag followed by the TEV recognition site (6xHis-FLAG-TEV-) with either the start of NPM1 or MCM2 coding sequence (see Table S2). Primers were used to amplify mouse NPM1 (Horizon discovery-Dharmacon) and MCM2, the product digested with BamH1 and EcoR1, and ligated with digested pFastbac1 (Invitrogen). Ligated product was transformed into XL10 Gold competent cells and colonies were picked and correct cloning confirmed by Sanger DNA sequencing.

For subcloning of GFP-ARF16-PB1-P2A-OsTIR1 into a TIGRE donor plasmid, GFP-ARF16-PB1-P2A-OsTIR-SV40(PolyA) was amplified from pMGS56, a gift from Michael Guertin (Addgene plasmid # 129668; http://n2t.net/addgene:129668; RRID:Addgene_129668), and digested with Mlu1 and Not1. The pEN396 TIGRE donor plasmid, a gift from Benoit Bruneau (Addgene plasmid # 92142; http://n2t.net/addgene:92142; RRID:Addgene_92142), was digested with Mlu1 and Not1 and ligated to digested GFP-ARF16-PB1-P2A-OsTIR-SV40(PolyA). The ligated pEN396-GFP-ARF16-PB1-P2A-OsTIR-SV40(PolyA) was transformed into XL10 Gold competent cells and correct cloning confirmed by Sanger DNA sequencing. The pX330-EN1201 (Cas9 + sgRNA against mouse TIGRE acceptor locus) was used along with pEN396-GFP-ARF16-PB1-P2A-OsTIR-SV40(PolyA) to make inducible AID-NPM1 KH2 mESCs, as described above.

For cloning of IAA17 (amino acids 71-113 of the AID protein) into pINTA-N3-dCas9-BirA (previously described (*1*), Gibson cloning was performed. The ~250 bp gBlock containing BirA-linker(5xAla-Gly)-AID-STOP (5′-CGACAAGCAGGGAGCTCTGCTGCTGGAGCAGGACGGAATCATCAAGCCCTGGATGGGCGGAGAAATCTCCCTGAGAAGCGCAGAGAAGGGAGCTGGTGCAGGCGCTGGAGCGGGTGCCCCTAAAGATCCAGCCAAACCTCCGGCCAAGGCACAAGTTGTGGGATGGC CACCGGTGAGATCATACCGGAAGAACGTGATGGTTTCCTGCCAAAAATCAAGCGGTGGCCCGGAGGCGGCGGCGTTCGTGTAG-3′) was Gibson cloned into pINTA-N3-dCas9-BirA using primers described in Table S2. Transformation of the Gibson reaction followed by Sanger DNA sequencing of clones provided confirmation of the correct fusion of dCas9-BirA-AID.

To generate pLVX-TRE3G-mCherry-HA-NPM1wt for rescue experiments as well as HA-NPM1AD1 and NPM1AD2 mutants, the restriction enzymes Mlu1 and EcoR1 were used to clone into the pLVX-TRE3G-mCherry (Clontech). For the full length amplification of NPM1, primers were synthesized from IDT with forward primer containing the Mlu1 restriction site, Kozak sequence, the HA-tag sequence, and a 5x Glycine linker (Table S2). Next, for each of the acidic mutants (HA-NPM1AD1, HA-NPM1AD2, and HA-NPM1AD1+AD2), PCR-fusion was conducted to delete 120-133bp and/or 159-188bp of NPM1 cDNA (Table S2). PCR products were then digested with Mlu1 and EcoR1 followed by ligation and transformation into XL10 Gold competent cells. Colonies were picked and correct cloning confirmed by Sanger DNA sequencing.

#### Protein purification using baculovirus expression system

The purification of PRC2 was described previously (*31*). To purify mouse 6xHis-FLAG-TEV-NPM1 and 6xHis-FLAG-TEV-MCM2, the proteins were expressed in Sf9 cells by baculovirus infection. After 60 hr of infection, Sf9 cells were resuspended in BC150 buffer (25 mM Hepes-NaOH, pH 7.8, 1 mM EDTA, 150 mM NaCl, 10 % glycerol, 1 mM DTT, and 0.1 % NP-40) with protease inhibitors (1 mM phenylmethlysulfonyl fluoride (PMSF), 0.1 mM benzamidine, 1.25 mg/ml leupeptin and 0.625 mg/ml pepstatin A) and phosphatase inhibitors (20 mM NaF and 1 mM Na3VO4). Cells were lysed by sonication (Fisher Sonic Dismembrator model 100), and NPM1 or MCM2 were tandemly purified through Ni-NTA agarose beads (Qiagen), FLAG-M2 agarose beads (Sigma), digested with His-TEV protease, and subjected to another purification with Ni-NTA agarose beads to remove any TEV contaminant as well as tagged full-length proteins (final product shown in Fig. S1D).

#### Whole cell extract and western blotting

Cells were harvested and lysed with CHAPS-Urea buffer (50 mM Tris-HCl, pH 7.9, 8 M Urea, and 1% CHAPS) containing protease inhibitors (0.2 mM PMSF, 1 μg/mL Pepstatin A, 1 μg/mL Leupeptin, and 1 μg/mL Aprotinin) and phosphatase inhibitors (10 mM NaF and 1 mM Na3VO4). The cell suspension was briefly sonicated (40% amplitude, 5 strokes) and centrifuged at 20,000 x g at 4°C for 20 min. The supernatant was collected and protein concentrations were quantified via a bicinchonic acid (BCA) assay. Proteins were separated using a 6%–12% BIS-TRIS SDS PAGE gel, and transferred onto a PVDF membrane. Membranes were blocked with 5% milk in PBST at RT for 1 hr and incubated with primary antibody overnight at 4°C. Membranes were washed 3 times with TBST and then incubated with HRP-conjugated secondary antibodies for 1 hr at RT, followed by exposure to ECL. Antibodies: NPM1 Abcam Cat# ab15440, H3K27Me3 Cell Signaling Cat# 9733, H3K27Me2 Cell Signaling Cat# 9728, H3 Abcam Cat# ab12079, H4 Abcam Cat# ab10158, Tubulin Abcam Cat# ab6046, Cas9 Millipore Cat# MAC133-clone7A9, EED in-house, EZH2 in-house, SUZ12 Cell Signaling Cat# 3737, H3K36me3 Abcam Cat# ab9050, and H3K9me3 Abcam Cat# ab8898.

#### *In vivo* biotinylation followed by native chromatin preparation

For *in vivo* biotinylation experiments using FLAG-BAP-H3.1; AID-NPM1 Kh2 mESCs, a similar protocol as described in (*1*) was followed. Briefly, mESCs were G1/S-synchronized with double thymidine (as described above). mESCs were given a 6 hr pulse of 2 μg/ml of doxycycline and 1 μg/ml of exogenous biotin during the latter half of the second thymidine treatment (12-13 hr). Next, mESCs were either harvested at the G1/2-block (starting time point, 0 hr) or washed with PBS and released to 2i media containing 2.5 mM Auxin for the time points indicated. For preparation of native MNase chromatin, cells were harvested using hypotonic lysis TMSD buffer (40 mM Tris, pH 7.5, 5 mM MgCl2, 0.25 M Sucrose with protease inhibitors), nuclei were then resuspended in NIB-250 (15 mM Tris-HCl, pH 7.5, 60 mM KCl, 15 mM NaCl, 5 mM MgCl2, 1 mM CaCl2, 250 mM Sucrose with protease inhibitors) containing 0.3% NP-40, and the chromosome pellet washed with NIB-250 buffer. After washes, the pellet was resuspended in MNase digestion buffer (10 mM Hepes, 50 mM NaCl, 5 mM MgCl2, 5 mM CaCl2 with protease inhibitors) and treated with MNase (Sigma) until a DNA fragment size of 150-300 bp (1-2 nucleosomes) was attained. The MNase reaction was stopped by addition of EGTA, spun down, and the supernatant was placed in a fresh tube. The chromatin pellet was further processed by adding the same volume of BC500 (40 mM Tris, pH 7.5, 5 mM MgCl2, 500 mM NaCl and 5% glycerol, and protease inhibitors) with EGTA, incubated for 30 min while rotating at 4°C, spun, and BC500 supernatant was pooled with an equal volume of MNase-treated supernatant to acquire the starting chromatin material for chromatin immunoprecipitations (IPs).

#### Native ChIP-qPCR

The protocol for native H3K27me3 and biotin chromatin IPs was described previously (*1*). Briefly, MNase-treated chromatin as above was pre-blocked with dynabeads protein G (Invitrogen), spun down, and the IP was set up with 100 μg of pre-cleared chromatin and 0.5 μg of *Drosophila* S2 chromatin (spike-in at a 1:100 concentration) in 10 μg of biotin antibody (Bethyl A150-109A), 4 μg of H3K27me3 (Cell Signaling C36B11), or 4 ug of H3K4me3 (Abcam ab8580) and 0.2 ug of H2AV (Active Motif Cat # 39715) and incubated overnight with slow rotation at 4°C. IPs were then washed three times with 1 ml BC300 buffer (40 mM Tris, pH 7.5, 5 mM MgCl2, 300 mM NaCl and 5% glycerol with protease inhibitors), once with 1 ml BC100 buffer (40 mM Tris, pH 7.5, 5 mM MgCl2, 100 NaCl and 5% glycerol with protease inhibitors), and a quick wash with 1 ml TE + 50 mM NaCl. Beads were then resuspended in 125 μl of TE, 3 μl of 10% SDS (TES) and incubated at 65°C for 1 hr followed by digestion for 2-4 hr with 8 μg of proteinase K at 55°C while shaking. All samples were PCR column purified, eluted in 50 μl, and diluted 1:4 with water for further qPCR studies.

For qPCR quantification, 5 μl SYBR Green I Master mix (Roche), ROX reference dye, 1 μl 5 μM primer pair, 4 μl-diluted DNA were mixed for PCR amplification, and detected by QuantStudio 5 real-tme PCR systems instrument (ThermoFisher Scientific). The data was then quantified and described in the corresponding figure legends. For *Drosophila* S2 chromatin, primers were FWD: TGGCTAGACTTTTGCGTCCT and REV:TACCAAAAGCCGTCCAAATC. For tiling of the *Gata2* and *Gata6* locus, qPCR primers were as previously listed in Escobar *et al*., 2019. Native biotin enrichment levels were normalized to 5% input followed by *Drosophila* chromatin spike-in levels. For time course experiments, data was minus-Dox (-Dox) control normalized and error bars represent standard deviation of three biological replicates. GraphPad Prism 7.0 was used for statistical analysis (2way ANOVA). A p value ≤ 0.05 was considered statistically significant; *** denotes p<0.001 and **** denotes p<0.0001.

#### ChIP and proteins digestion

The Rapid immunoprecipitation mass spectrometry of endogenous proteins (RIME) protocol (*32*), was used to identify protein-nucleosome interactions. Briefly, FLAG-BAP-H3.1 KH2 mESCs expressing HA-MCM2 and KH2 mESCs (untagged controls) were crosslinked with 1% (vol/vol) formaldehyde for 4 min at RT, quenched with glycine and chromatin isolated with nuclear extraction buffer 1 (LB1), pelleted and resuspended in LB2. Chromatin was pelleted and resuspended in LB3 and sheared to 200-600 bp using Bioruptor (Diagnode). Cleared chromatin lysate samples were subjected to tandem purification, as follows. First, H3.1 chromatin was IP’ed using anti-FLAG M2 affinity gel (Sigma-Aldrich) overnight at 4°C. H3.1-enriched protein complexes were eluted twice with BC100 containing FLAG peptide (Sigma-Aldrich) (first elution for 6 hr and second elution for 2 hr), followed by a second IP using antibodies against either H3K27me2/3 or HA (Abcam Cat# ab9110) overnight. Dynabeads ProteinG beads (Invitrogen) were then added the next day for 2-4 hrs and the ChIP samples were washed ten times with 1ml of RIPA buffer. All buffers included protease and phosphatase inhibitors (0.2 mM PMSF, 1 μg/mL Pepstatin A, 1 μg/mL Leupeptin, and 1 μg/mL Aprotinin) and phosphatase inhibitors (10 mM NaF and 1 mM Na3VO4), as well as 5 mM sodium butyrate. After the last RIPA wash and prior to trypsin digestion, beads were washed with 100 mM ammonium bicarbonate.To digest proteins, beads were resuspended in 50 mM ammonium bicarbonate containing 20ng/μl trypsin/Lys-C mix (Promega) followed by overnight incubation at 37°C with vigorous shaking. Reactions were then transferred to new tubes, acidified by mixing with 20%heptafluorobutyric acid added to 1% final concentration followed by 5-min incubation at room temperature and 5-min centrifugation at 16000×g. Peptides from clarified samples were desalted using Pierce C18 spin tips (Thermo Scientific) according to manufacturer instructions, dried under vacuum and dissolved in 0.1% formic acid. Peptides concentration was measured on Nanodrop One (Thermo Scientific) at 205 nm.

#### LC-MS analysis

Peptides were analyzed on Orbitrap Lumos Fusion mass spectrometer coupled with Dionex Ultimate 3000 UHPLC (Thermo Scientific). 1-2 μg of peptides were resolved on 50-cm long EASY-Spray column (Thermo Scientific) at flow rate 0.2 μl/min over 90-min gradient 4-40% acetonitrile in 0.1% formic acid followed by steep 5-min increase to 96% acetonitrile and 5-min step elution with 96% acetonitrile. Data-dependent acquisition method was based on published protocol (*33*) except each cycle was set to last 2 s.

#### Mass spectrometry data analysis

Raw mass spectrometry data were processed with Proteome Discoverer 2.1.1.21 (Thermo Scientific) and/or MaxQuant 1.6.6.0 (*34, 35*). Protein database supplied to programs comprised of mouse proteome (https://www.uniprot.org/proteomes/UP000000589) combined with list of known protein contaminants (distributed with MaxQuant). Sequest HT search engine within Proteome Discoverer was run with default mass tolerance parameters. Variable modifications were: methionine oxidation, cysteine carbamidomethylation, lysine and protein N terminus acetylation, lysine and arginine methylation, and phosphorylation on serine, threonine, and tyrosine. MS1-based label-free quantitation was done using Precursor Ions Area Detector module in Proteome Discoverer. Individual peptide intensities were normalized to total peptide amount. Protein intensities were calculated using Top N method that considered three top-most abundant peptides per protein (*36*). In MaxQuant, Andromeda search engine was run with default parameters for mass tolerance. Variable modifications were: methionine oxidation, cysteine carbamidomethylation, and protein N terminus acetylation. To increase number of identified and quantitated peptides, match between runs option was enabled. For label-free quantitation, integrated intensity was used instead of maximum intensity. In further analyses, protein intensities were normalized to that of Protein G followed by subtracting intensity in untagged controls for each dataset.

#### RNA purification, RT-qPCR, and RNA-seq

Total RNA extractions were performed using Roche High pure RNA isolation kit. Superscript III Reverse Transcription Reagents (Invitrogen) and random hexamers were used to prepare cDNAs. For qPCR quantification, 5 μl SYBR Green I Master mix (Roche), ROX reference dye, 1 μl IDT PrimeTime Primer set for corresponding assays, 1 μl cDNA were mixed for PCR amplification, and detected by QuantStudio 5 real-time PCR systems instrument. For quantitative analysis, Gata2 and Gata6 expression was normalized to Actb expression. For RNA-seq, the first strand was synthesized using reverse transcription via Superscript III and random hexamers. The second strand was synthesized with dUTP to generate strand asymmetry using DNA Pol I (NEB, M0209L) and the *E. coli ligase* (Enzymatics, L6090L). RNA-seq libraries were constructed using the protocol described in Escobar et al. 2019, quantified by Qubit dsDNA HS Assay Kitquality, and checked by High Sensitivity D1000 ScreenTape. Libraries were then sequenced as 50 bp single-end reads on NovaSeq 6000 platform.

#### RNA-seq analysis

Reads were aligned to the mouse reference genome mm10 using STAR with parameters: -- outFilterMismatchNoverLmax 0.2 --outFilterMultimapNmax 1 --outSAMstrandField intronMotif --outSAMmapqUnique 60 --twopassMode Basic --outSJfilterReads Unique -- outFilterIntronMotifs RemoveNoncanonical. Gene counts were calculated using featureCounts with parameters: -p -s 2 -t exon, and RefSeq mm10 annotation downloaded from GENCODE. The output gene count tables were used as input into DeSeq2 for normalization and differential expression analysis. Bigwig files were generated using deeptools and default parameters for visualization in IGV.

#### ChIP-seq data analysis

Reads were aligned to the mouse reference genome mm10 and dm6 for spike-in samples, using Bowtie2 with default parameters. Reads of quality score less than 30 were removed using samtools and PCR duplicates were marked using picard. Regions in mm10 genome blacklist was removed using bedtools and bigwig files were generated using deeptools and parameters: --binSize 50 -- normalizeUsing RPKM --ignoreDuplicates --ignoreForNormalization chrX --extendReads 250 for visualization in IGV. Peaks were called using MACS2 with parameters: -f BAM -g mm --keep-dup all --broad --broad-cutoff 0.1. Genomic peak annotation was performed with the R package ChIPseeker considering the region ± 3 kb around the TSS as the promoter. Peak overlapping analysis was performed using the Python package Intervene and visualized using the Python package Matplotlib.

For visualization of ChIP-seq, uniquely aligned reads mapping to the mouse genome were normalized using dm6 spike-in as described previously (*37*). Heatmaps were performed using the functions computeMatrix followed by plotHeatmap and plotProfile from deepTools.

#### CUT&RUN library preparation and data analysis

The CUT&RUN experiments were performed using the CUTANA™CUT&RUN assay (EpiCypher) under the manufacturer’s instructions. Briefly, the nuclei of cells were isolated using the TMSD buffer as previously described (*38*). The nuclei were bound by activated Concanavalin A beads, followed by nuclear membrane permeabilization, incubation of primary antibody overnight, and binding of antibody to pA-MNase fusion protein. The MNase was activated by CaCl at 4° C for 1 hr. The DNA released into the supernatant were purified and subjected to library preparation and next-generation sequencing (NovaSeq 6000, Illumina). Data were analyzed using the same pipeline as previously described above for ChIP-seq,

**Figure S1.**
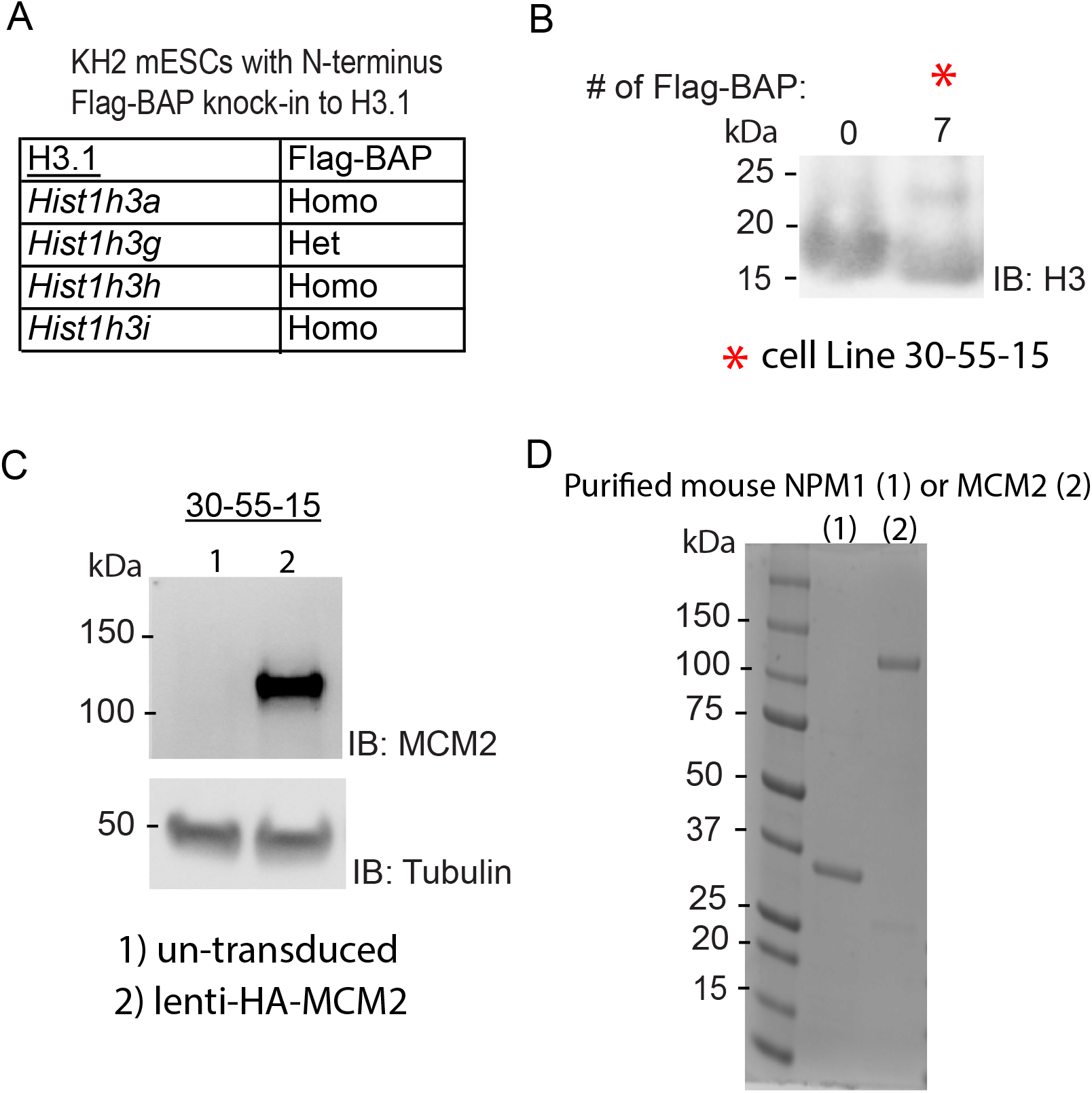
Characterization of 30-55-15 cells and protein expression. **(A)** Overview and genotype of KH2 mESC lines containing FLAG-BAP knockins to four H3.1 loci and **(B)** western blot analysis of acid-extracted histone H3 using anti-H3 antibody. The upper band corresponds to the endogenous number of FLAG-BAP-H3 copies in the cell and the red asterisk denotes the cell line (30-55-15) used for further studies. **(C)** Western blot of HA-tagged MCM2 expressed from lentivirus after infection of 30-55-15 cells, used for proteomic studies. **(D)** SDS-PAGE analysis of purified, recombinant NPM1 and MCM2 visualized with Coomassie blue staining. Molecular weight markers are shown.

**Figure S2.**
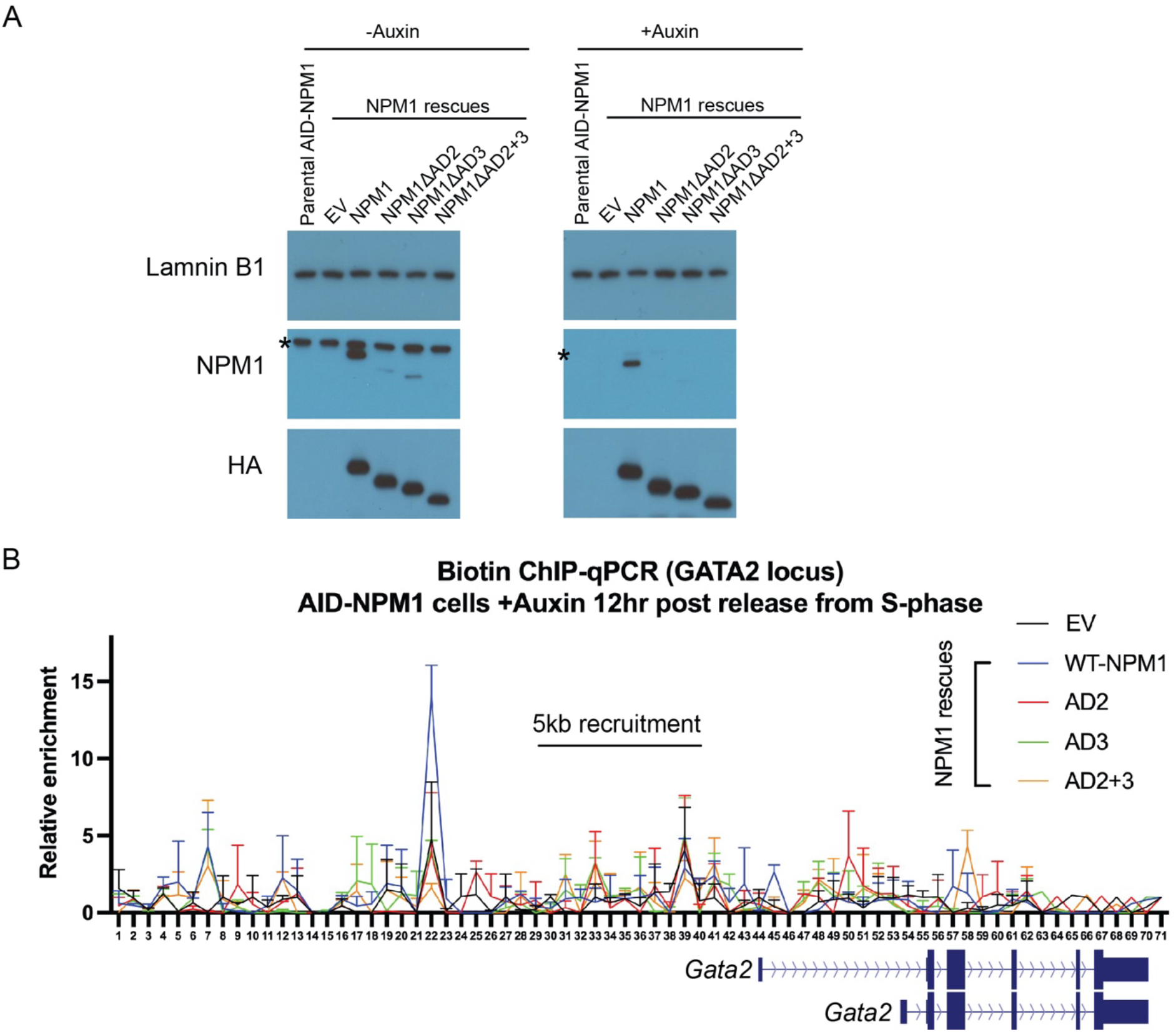
NPM1-mediated parental histone inheritance requires its acidic patches involved in its chaperone activity. **(A)** Western blot of Laminin B1, NPM1, and HA tag for the AID-NPM1 cells rescued with empty vector (EV), HA-tagged wild type NPM1, or HA-tagged NPM1 acidic patch mutants (AD2, AD3, or AD2+3) without auxin (left) or with auxin treatment (right) for 12 hr upon release into S-phase, as depicted in Fig 4I. Asterisk indicates the expected size of AID-NPM1. **(B)** Biotin ChIP-qPCR using a tiling array of primers flanking the dCas9-BirA-AID recruitment site in the cell lines denoted in **(A)**, right panel.

## References and Notes

1. T. M. Escobar et al., Active and Repressed Chromatin Domains Exhibit Distinct Nucleosome Segregation during DNA Replication. Cell 179, 953–963 e911 (2019).

2. G. Schlissel, J. Rine, The nucleosome core particle remembers its position through DNA replication and RNA transcription. Proc Natl Acad Sci U S A 116, 20605–20611 (2019).

3. T. M. Escobar, A. Loyola, D. Reinberg, Parental nucleosome segregation and the inheritance of cellular identity. Nat Rev Genet 22, 379–392 (2021).

4. R. Margueron, D. Reinberg, The Polycomb complex PRC2 and its mark in life. Nature 469, 343–349 (2011).

5. J. A. Simon, R. E. Kingston, Occupying chromatin: Polycomb mechanisms for getting to genomic targets, stopping transcriptional traffic, and staying put. Mol Cell 49, 808–824 (2013).

6. L. Di Croce, K. Helin, Transcriptional regulation by Polycomb group proteins. Nat Struct Mol Biol 20, 1147–1155 (2013).

7. S. Aranda, G. Mas, L. Di Croce, Regulation of gene transcription by Polycomb proteins. Sci Adv 1, e1500737 (2015).

8. J. R. Yu, C. H. Lee, O. Oksuz, J. M. Stafford, D. Reinberg, PRC2 is high maintenance. Genes Dev 33, 903–935 (2019).

9. R. Margueron et al., Role of the polycomb protein EED in the propagation of repressive histone marks. Nature 461, 762–767 (2009).

10. O. Oksuz et al., Capturing the Onset of PRC2-Mediated Repressive Domain Formation. Mol Cell 70, 1149–1162 e1145 (2018).

11. N. Rhind, D. M. Gilbert, DNA replication timing. Cold Spring Harb Perspect Biol 5, a010132 (2013).

12. A. Groth et al., Regulation of replication fork progression through histone supply and demand. Science 318, 1928–1931 (2007).

13. N. Richet et al., Structural insight into how the human helicase subunit MCM2 may act as a histone chaperone together with ASF1 at the replication fork. Nucleic Acids Res 43, 1905–1917 (2015).

14. J. K. Box et al., Nucleophosmin: from structure and function to disease development. BMC Mol Biol 17, 19 (2016).

15. L. J. Frehlick, J. M. Eirin-Lopez, J. Ausio, New insights into the nucleophosmin/nucleoplasmin family of nuclear chaperones. Bioessays 29, 49–59 (2007).

16. M. Okuwaki, K. Matsumoto, M. Tsujimoto, K. Nagata, Function of nucleophosmin/B23, a nucleolar acidic protein, as a histone chaperone. FEBS Lett 506, 272–276 (2001).

17. M. Okuwaki, A. Iwamatsu, M. Tsujimoto, K. Nagata, Identification of nucleophosmin/B23, an acidic nucleolar protein, as a stimulatory factor for in vitro replication of adenovirus DNA complexed with viral basic core proteins. J Mol Biol 311, 41–55 (2001).

18. S. Grisendi et al., Role of nucleophosmin in embryonic development and tumorigenesis. Nature 437, 147–153 (2005).

19. K. Nishimura, T. Fukagawa, H. Takisawa, T. Kakimoto, M. Kanemaki, An auxin-based degron system for the rapid depletion of proteins in nonplant cells. Nat Methods 6, 917–922 (2009).

20. K. M. Sathyan et al., An improved auxin-inducible degron system preserves native protein levels and enables rapid and specific protein depletion. Genes Dev 33, 1441–1455 (2019).

21. B. Falini et al., Cytoplasmic nucleophosmin in acute myelogenous leukemia with a normal karyotype. N Engl J Med 352, 254–266 (2005).

22. L. Brunetti et al., Mutant NPM1 Maintains the Leukemic State through HOX Expression. Cancer Cell 34, 499–512 e499 (2018).

23. V. Azuara et al., Chromatin signatures of pluripotent cell lines. Nat Cell Biol 8, 532–538 (2006).

24. B. E. Bernstein et al., A bivalent chromatin structure marks key developmental genes in embryonic stem cells. Cell 125, 315–326 (2006).

25. T. S. Mikkelsen et al., Genome-wide maps of chromatin state in pluripotent and lineage-committed cells. Nature 448, 553–560 (2007).

26. P. Voigt et al., Asymmetrically modified nucleosomes. Cell 151, 181–193 (2012).

27. K. H. Hansen et al., A model for transmission of the H3K27me3 epigenetic mark. Nat Cell Biol 10, 1291–1300 (2008).

28. C. Alabert et al., Nascent chromatin capture proteomics determines chromatin dynamics during DNA replication and identifies unknown fork components. Nat Cell Biol 16, 281–293 (2014).

29. A. Hugues, C. S. Jacobs, F. Roudier, Mitotic Inheritance of PRC2-Mediated Silencing: Mechanistic Insights and Developmental Perspectives. Front Plant Sci 11, 262 (2020).

30. M. Okuwaki, The structure and functions of NPM1/Nucleophsmin/B23, a multifunctional nucleolar acidic protein. J Biochem 143, 441–448 (2008).

31. C. H. Lee et al., Allosteric Activation Dictates PRC2 Activity Independent of Its Recruitment to Chromatin. Mol Cell 70, 422–434 e426 (2018).

32. H. Mohammed et al., Rapid immunoprecipitation mass spectrometry of endogenous proteins (RIME) for analysis of chromatin complexes. Nat Protoc 11, 316–326 (2016).

33. S. Davis et al., Expanding Proteome Coverage with CHarge Ordered Parallel Ion aNalysis (CHOPIN) Combined with Broad Specificity Proteolysis. J Proteome Res 16, 1288–1299 (2017).

34. J. Cox, M. Mann, MaxQuant enables high peptide identification rates, individualized p.p.b.-range mass accuracies and proteome-wide protein quantification. Nat Biotechnol 26, 1367–1372 (2008).

35. S. Tyanova, T. Temu, J. Cox, The MaxQuant computational platform for mass spectrometry-based shotgun proteomics. Nat Protoc 11, 2301–2319 (2016).

36. J. C. Silva, M. V. Gorenstein, G. Z. Li, J. P. Vissers, S. J. Geromanos, Absolute quantification of proteins by LCMSE: a virtue of parallel MS acquisition. Mol Cell Proteomics 5, 144–156 (2006).

37. D. A. Orlando et al., Quantitative ChIP-Seq normalization reveals global modulation of the epigenome. Cell Rep 9, 1163–1170 (2014).

38. N. D. Fogarty, K. L. Marhaver, Coral spawning, unsynchronized. Science 365, 987–988 (2019).

